# Direct Repeats Co-occur with Few Short Dispersed Repeats in Plastid Genome of A Spikemoss, *Selaginella vardei* (Selaginellaceae, Lycophyta)

**DOI:** 10.1101/505842

**Authors:** Hong-Rui Zhang, Xian-Chun Zhang, Qiao-Ping Xiang

## Abstract

**Background:** It is hypothesized that the highly conserved inverted repeat (IR) structure of land plant plastid genomes (plastomes) is beneficial for stabilizing plastome organizations, whereas the mechanism of the occurrence and stability maintenance of the newly reported direct repeats (DR) structure was yet awaiting further exploration. Here we introduced the DR structure of plastome in *Selaginella vardei* (Selaginellaceae, Lycophyta), trying to elucidate the mechanism of DR occurrence and stability maintenance.

**Results:** The plastome of *S. vardei* is 121,254 bp in length and encodes 76 different genes, of which 62 encode proteins, 10 encode tRNAs and four encode rRNAs. Unexpectedly, the two identical rRNA gene regions (13,893 bp) are arranged into DR, and a ca. 50-kb *trn*N-*trn*F inversion spanning one DR copy exists in *S. vardei*, comparing to the typical IR organization of *Isoetes flaccida* (Isoetaceae, Lycophyta). We find extremely rare short dispersed repeats (SDRs) in plastome of *S. vardei* and is confirmed in its closely related species *S. indica*. The occurrence time of DR in Selaginellaceae is estimated at late Triassic (ca. 215 Ma) based on the phylogenetic framework of land plants.

**Conclusions:** We propose that the unconventional DR structure, co-occurred with extremely few SDRs, plays key role in maintaining the stability of plastome, and reflects a relic of the environmental upheaval during extinction event. We suggest that the ca. 50-kb inversion resulted in the DR structure, and recombination between DR regions is confirmed to generate multipartite subgenomes and diverse multimers, which shed lights on the diverse structures in plastome of land plants.

## Background

Chloroplasts play a crucial role in maintaining life on earth through the process of photosynthesis in plants [1]. Chloroplasts in land plants have genomes that are widely conserved due to the constantly high selective pressures of photosynthesis [2]. Most plastomes are characterized by a quadripartite structure, which comprises two copies of an inverted repeat (IR) separating the large (LSC) and small (SSC) single copy regions. The size of land plant plastomes usually ranges from 108 to 165 kb, and they generally contain 110-130 distinct genes including about 30 transfer RNA (tRNA) genes, four ribosomal RNA (rRNA) genes, and approximately 80 protein-coding genes involved in photosynthesis or other metabolic processes [3, 4]. The typical IR is usually 20 – 30 kb and the genes that form the core of the IR encode the ribosomal RNAs (23S, 16S, 5S and 4.5S) [5]. While plastomes of most land plants possess the typical IR structure, several lineages of land plants only retain one copy of the IR, such as in *Carnegiea gigantean* (Cactaceae) [6], *Erodium* (Geraniaceae) [7], and an IR-lacking clade of Fabaceae [8], as well as conifers [9], in which one IR has been either extremely shortened or completely lost.

The conserved IR structure across land plants was hypothesized to function as stabilizing the entire plastome against major sequence rearrangements [10]. JD Palmer and WF Thompson [11] showed that rearrangement events were extremely rare in genomes with an IR, but increased remarkably in frequency when one IR copy was absent. Their hypothesis is supported by some recent studies using next generation sequencing. For example, *Trifolium* [12] and *Erodium texanum* [13] lack one copy of the IR and have highly rearranged plastomes. However, plastomes of *Pelargonium* [14] and *Trachelium* [15] are also highly rearranged, despite the presence of the IR structure, so that the existence of the IR appears to be not sufficient to stabilize the plastome structure [16]. A related hypothesis suggests that the incidence of short dispersed repeats (SDRs) is correlated with instability of plastomes. Extensive studies have shown that genomes with massive rearrangement events tend to contain a high frequency of SDRs (*Trifolium* [12]; *Trachelium* [15]; *Pelargonium* [14]), whereas genomes containing virtually no SDRs have the conserved organization (*Erodium* [13]; algae [17, 18]).

Algal plastomes have shown extensive variation in size and structure compared to land plants [19–21]. Certain dinoflagellates contain several 2 to 3 kb minicircles mostly encoding only a single gene [22], whereas Cladophorales algae possess the fragmented linear ssDNA molecules folding into hairpin configurations [23]. The number and orientation of rRNA encoding repeats are also much more variable in algae than in land plants [24]. One to five copies of rRNA encoding repeat are tandemly arranged in *Euglena* (green algae) [25], whereas in *Porphyra purpurea* (red alga), the two copies of rRNA encoding region are arranged into a direct repeat (DR) [26]. The mechanism of creating and maintaining this diversity remains unknown.

Among early diverging land plant lineages, *Selaginella tamariscina* (Selaginellaceae), has been recently reported to possess plastome with DR structure based on assembly from PacBio data [27]. However, the potential mechanism for its origin and unusual organization was left unknown and not discussed. Here we confirmed plastomes with DR structure in *Selaginella* subg. *Rupestrae*, (Selaginellaceae, *sensu* Weststrand and Korall (2016)). We discovered that the two copies of DR were probably caused by a ca. 50-kb *trn*F-*trn*N inversion that containing one DR copy. We also found extremely rare short SDRs in plastomes with DR comparing to that with IR in lycophyte. Given the subgenomes generated by the recombination between DR regions, we predict that the DR structure, co-occurred with extremely few SDRs, plays an important role in maintaining the stability of plastome and survived over 400 million years’ evolutionary history [28].

## Results

### The Unconventional Structure of the *S. vardei* Plastome

We sequenced the plastome of *S. vardei* (Fig. 1; MG272482) using Illumina HiSeq 2500 sequencing and assembled reads using *de novo* assembly methods. We combined the assembled contigs in Geneious 9.1.4 (Biomatters, Inc., Auckland, New Zealand; https://www.geneious.com) to extend them into one unit-genome. Reads coverage of the *S. vardei* plastome sequence is shown in Additional file 1, Fig. S1**c**. We also performed the assembly of the contigs in Bandage 0.8.1 [29], which similarly resulted in a single unit, circular plastome (Additional file 1, Fig. S1**d**). The plastome of *S. indica* (MK156801) was also assembled into a complete unit genome with DR structure (data not shown). The plastome organization and gene content are basically consistent with *S. vardei*, therefore, we, here, only described the detailed plastome information of *S. vardei*.

The plastome of *S. vardei* was 121,254 bp in length and contained 76 different genes with 62 protein coding genes, four rRNA genes, and 10 tRNA genes (Additional file 6, Table S1). Thus, the plastome size of *S. vardei* was much smaller than that of the published plastomes of *S. uncinata* and *S. moellendorffii*, apparently owing to gene losses, such as, of all 11 *ndh* genes and two more tRNA genes (*trn*Q and *trn*R). The plastome of *S. vardei* was similar to most land plants in having a quadripartite structure, separated by two copies of large rRNA encoding repeat (13,893 bp). However, the two copies of repeat were arranged into DR, with two single copy regions of almost equal size (47,676 bp and 45,792 bp, respectively).

**Fig. 1.**
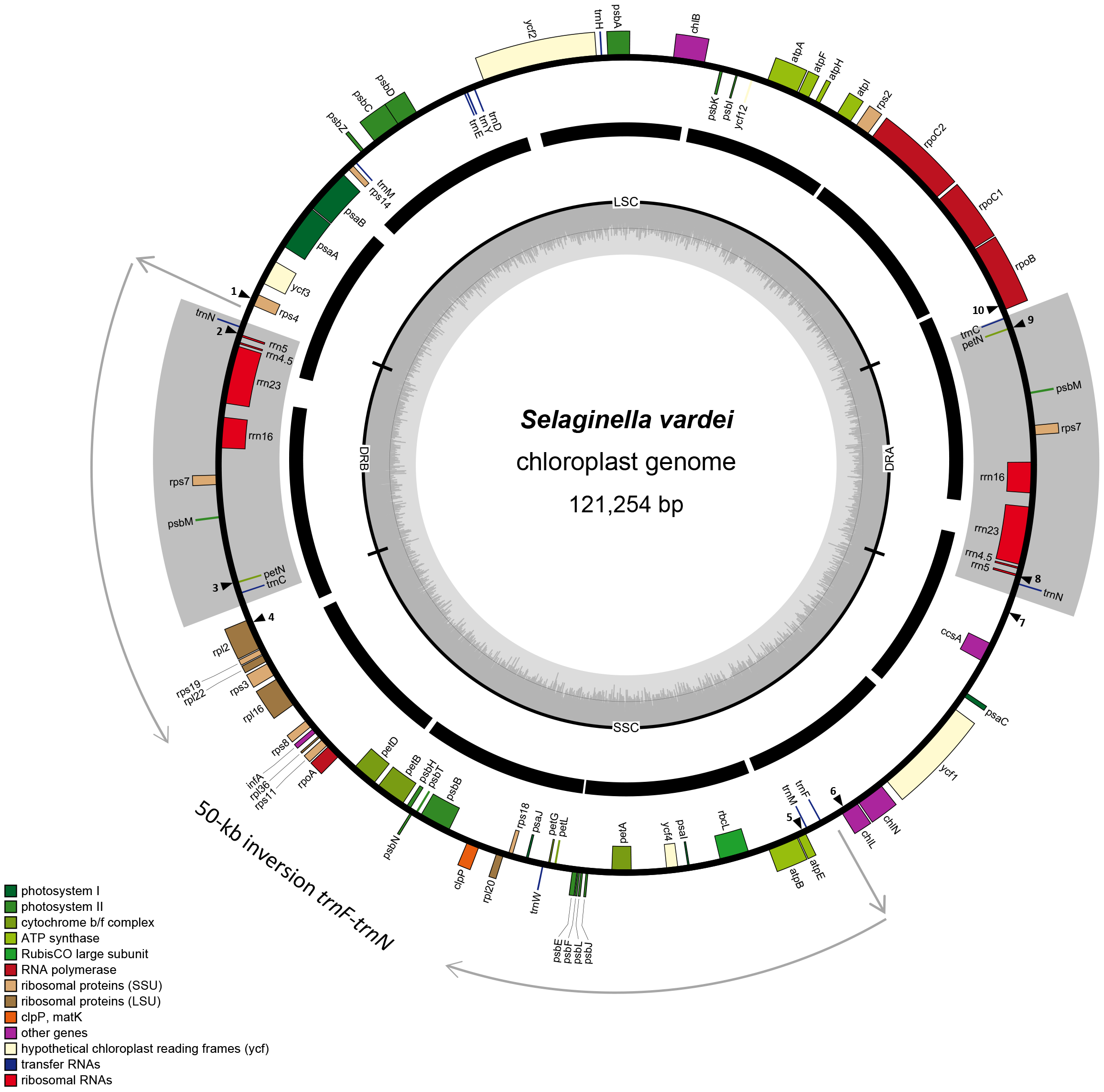
Unit-genome map of *S. vardei* shown as circular. The middle ring shows the mapping results of PCR data with assembled sequence. The arrows point to the endpoints of the ca. 50-kb fragment (from *trn*N to *trn*F) inversion. The black triangular and numbers showed the position of primers for confirming the boundaries of DR regions and the 50-kb inversion.

### Confirmation of the Plastome Structure of *S. vardei*

To verify the DR structure in *S. vardei* assembled with next generation data, we performed long PCR experiments across the whole plastome and obtained the sequences using Sanger sequencing methodology. The long PCR experiments resulted in 12 products of the expected length (ca. 7-11 kb) (Additional file 1, Fig. S1**a**) using 12 pairs of primers (Additional file 7, Table S2) and newly designed internal primers (Additional file 8, Table S3). We mapped the long PCR products to the assembled plastome of *S. vardei* (Fig. 1). All the intergenic regions were covered by Sanger sequencing.

In addition, the PCR product of the twelve pairs of primers at the boundaries of DR region and the ca. 50-kb inversion also supported the plastome structure of *S. vardei* (Additional file 1, Fig. S1**b**). The amplified fragments were consistent with the DR structure (1-2: *rps*4-*rrn*5 and 3-4: *pet*N – *rpl*2; 7-8: *ccs*A-*rrn*5 and 9-10: *pet*N-*rpo*B) and the ca. 50-kb inversion (1-2: *rps*4-*rrn*5 and 5-6: *atp*E-*chl*L) in *S. vardei*. Accordingly, a negative control consistent with amplification of a typical IR structure (1-3: *rps*4-*pet*N and 2-4: *rrn*5-*rpl*2; 7-9: *ccs*A-*pet*N and 8-10: *rrn*5-*rpo*B) and without the ca. 50-kb inversion structure (1-5: *rps*4-*atp*E and 2-6: *rrn*5-*chl*L) yielded no PCR product. We further checked the sequences at boundaries of the DR region and the ca. 50-kb inversion. The assembled sequences from NGS were identical with sequences from Sanger sequencing. The congruent results from Sanger with next generation sequencing strongly support the DR structure and the ca. 50-kb inversion in the *S. vardei* plastome.

### PCR Confirmation of DR Structure in Representatives of subg. *Rupestrae*

We performed PCR experiments on one additional individual of *S. vardei* and two other species representing *Selagenilla* subg. *Rupestrae* (Table 1, additional file 9, Table S4) and these yielded the expected products with consistent length (Additional file 2, Fig. S2**a**). PCR products were consistent with DR structure (1-2: *rps*4-*rrn*5 and 3-4: *pet*N-**rpl*2;* 7-8: *ccs*A-*rrn*5 and 9-10: *pet*N-*rpo*B) and the ca. 50-kb inversion (1-2: *rps*4-*rrn*5 and 5-6: *atp*E-*chl*L). The alignment sequences of the four representatives showed several insertion/deletions (Additional file 2, Fig. S2**b**). Sequence divergence is remarkable between *S. dregei* and the other two species. For example, a 475 bp deletion existed at the region of *ccs*A-*rrn*5 in the plastome of *S. dregei*, (collected from Kenya, Africa), which also exhibits a smaller sized PCR product than the other species (Additional file 2, Fig. S2**b**: 26698, 7-8).

**Table 1.** Representatives related for confirmation of plastome structure of *S. vardei*

| Species | Voucher | Locality |
| --- | --- | --- |
| *S. vardei* | Zhang X. C. et al. 836 (PE) | Tibet, China |
| *S. indica* | Zhang X. C. 5868 (PE) | Sichuan Province, China |
| *S. indica* | Zhang X. C. 6255 (PE) | Yunnan Province, China |
| *S. dregei* | Liu B. 26698 (PE) | Kenya |

### Gene Order of Plastomes in *S. vardei* and Other Lycophytes

Dot plot analysis (Additional file 3, Fig. S3) showed that the gene order of the plastome of the *S. vardei* was considerably divergent from other species of lycophytes.

Plastomes of *Huperzia serrata* and *Isoetes flaccida* were basically syntenic representing the ancestral organization for lycophytes (Additional file 3, Fig. S3**a**). However, plastomes of *S. vardei* were quite divergent when comparing with its sister group *I. flaccida* (Additional file 3, Fig. S3**b**). A large inversion was present in *S. vardei* that encompasses a ca. 50-kb region from *trn*N to *trn*F, its endpoints lying between *rps*4 and *trn*N at one end and between *trn*F and *chl*L at the other (Figs. 1 and 2). In *I. flaccida*, two tRNA genes, *trn*L-UAA and *trn*T-UGU were situated between *rps*4 and *trn*F in LSC region, whereas *trn*N was at the border of the IRb/SSC adjacent to *ycf*2. In *S. vardei*, however, *trn*L-UAA and *trn*T-UGU were absent, *trn*F was located in the SSC, and *trn*N was located at the border of DRb/LSC next to *rps*4. Thus, we infer that the DR in *S. vardei* was caused by this ca. 50-kb inversion owing to one copy of the DR (*trn*N-*trn*C) located inside the inversion. In addition, *S. vardei* has an expanded DR that has duplicated psɓM through *trnC* from the LSC, and *rpo*B was located at the start position of LSC region. Gene *ycf*2 relocated to LSC, and chlL/chlN relocated to be adjacent to *ycf*1, which was consistent with the plastome of *H. serrata*.

**Fig. 2.**
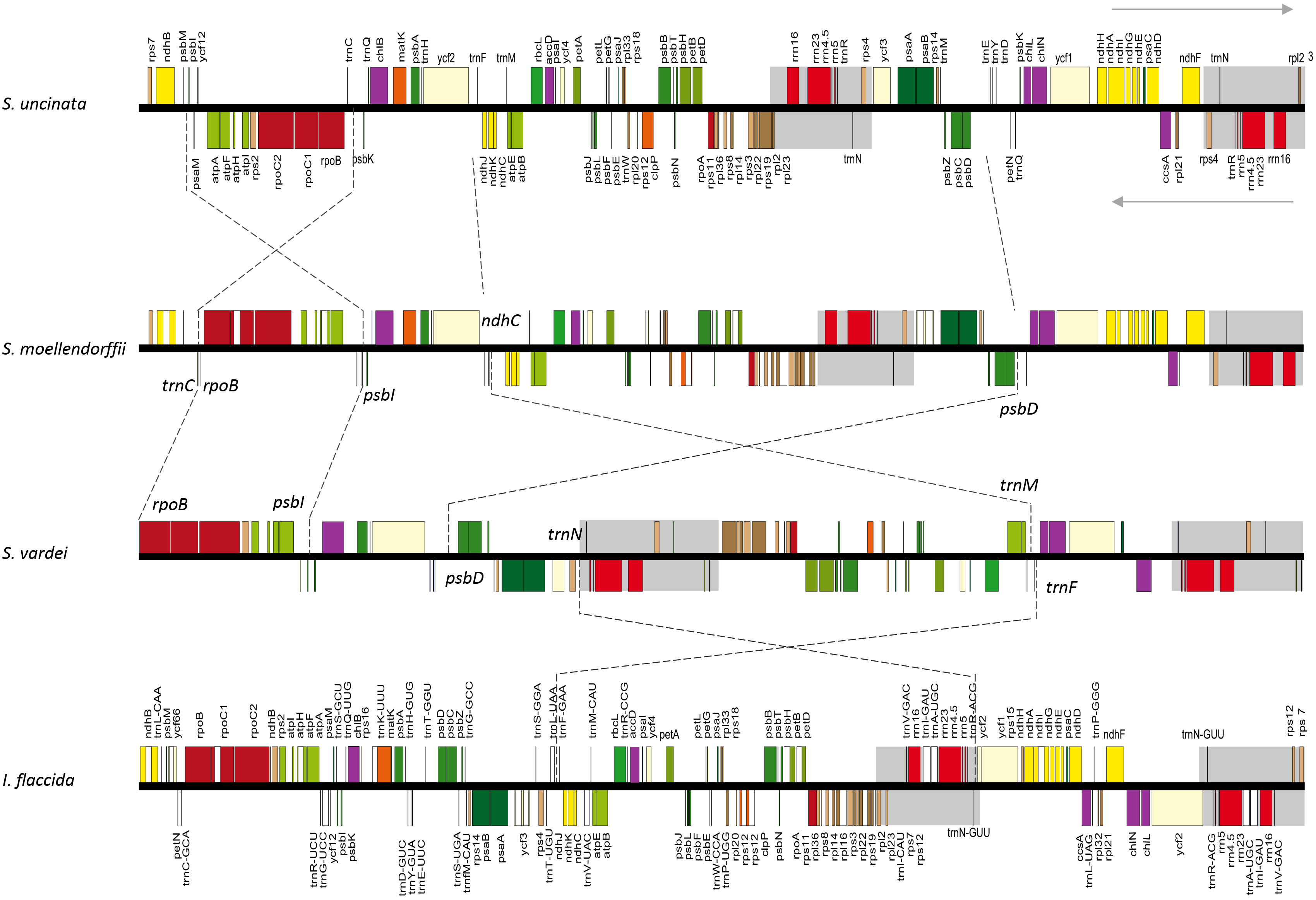
Linear maps of plastomes between *S. vardei* and other lycophytes showing the inversions. Upper arrow indicates genes in forward direction; lower arrow indicates genes in reverse direction.

One remarkable difference between the plastomes of *S. uncinata* and *S. moellendorffii* was that *S. moellendorffii* lacked a ca. 20-kb inversion (from *trn*C to *psb*I) existing in *S. uncinata* (Fig. 2). This inversion was also absent from the plastome of *S. vardei*, which suggests that the absence of this *trn*C-*psb*I inversion might be the ancestral state in Selaginellaceae. The previously published plastomes of *S. uncinata* [30] and *S. moellendorffii* [31] possess the typical IR structure, which was universal in other land plants [2]. Comparing the plastomes among *S. vardei* and the two published *Selaginella* species, our dot plot results showed that an inversion of the ca. 60-kb psbD-*trnM/ndhJ* (lost in *S. vardei*) fragment existed between the plastomes of *S. vardei* and *S. moellendorffii* (Additional file 3, Fig. S3**c**), whereas a ca. 65-kb inversion of *trnD-trnF* existed in plastomes between *S. vardei* and *S. uncinata* (Additional file 3, Fig. S3**d**).

### Short Dispersed Repeats (SDRs) in Plastomes of *S. vardei* and Other Species

Repeats analyses showed that SDRs in the Selaginellaceae plastomes with DR ((Fig. 3e, f) were obviously fewer than those with IR in other lycophytes (Fig. 3a-d and Additional file 10, Table S5). *Huperzia serrata* had the most SDRs with thirty-one whereas *S. vardei* and *S. indica* contained the fewest with only six and five, respectively. Furthermore, no SDRs were found at the endpoints of the inversion and the DR regions in the plastome of *S. vardei* and *S. indica*, and the length of all the SDRs in them was less than 50 bp. In *S. uncinata*, a pair of short repeats (46 bp) was located between the *psb*I and the *trn*C-*psb*K intergenic region, flanking the ca. 20-kb inversion from *psb*I-*trn*C (Fig. 3c and Additional file 3, Fig. S3**d**). Another pair of 98 bp short repeats located between *pet*A-*psb*J/*rpl*20-*psb*B in plastome of *S. uncinata* could also potentially cause inversions (Additional file 10, Table S5-3).

**Fig. 3.**
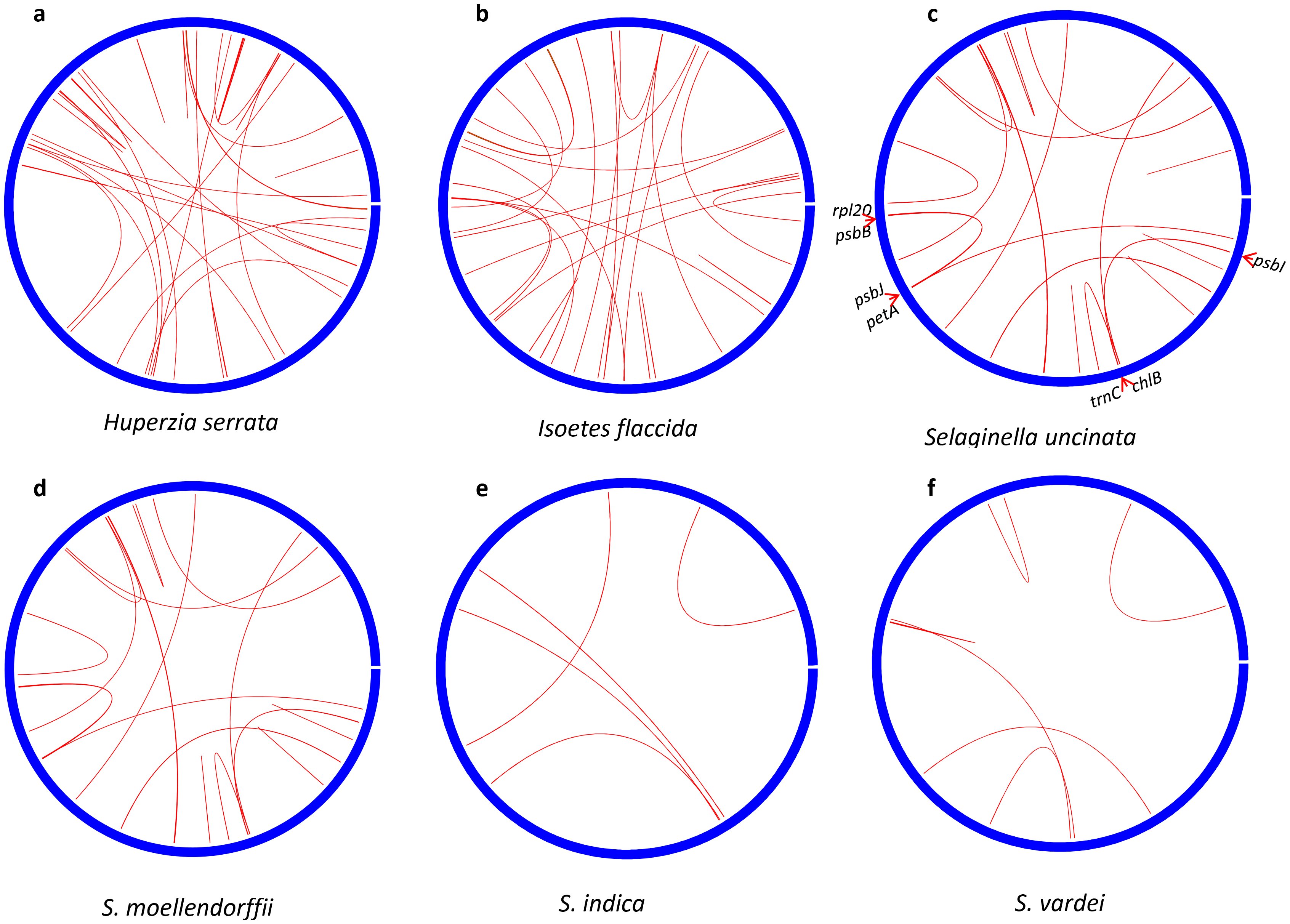
Repeats of *S. vardei* and other species in Lycophytes. The gap of each circle represents the start position and the direction is clockwise.

### The Evolution and Divergence Time of DR in Plastomes of Land Plants

Phylogenetic relationships based on 32 protein-coding genes of plastomes (Fig. 4) showed that subg. *Rupestrae* containing *S. vardei* and *S. indica* was early diverging compared to *S. tamariscina*, which was sister to the clade containing *S. moellendorffii* and *S. uncinata* in Selaginellaceae (Fig. 4). The split between Selaginellaceae and its sister family Isoetaceae was ca. 381 million years ago (Ma) [32]. We inferred the divergence time of subg. *Rupestrae* was at late Triassic (ca. 215 Ma) (Fig. 4). When we mapped the simplified structure of plastomes to the phylogenetic tree, it is clear that most plastomes of land plants possessed the typical IR structure. In plastome of ferns, the rRNA genes were arranged in reverse order in Schizaeales and core leptosporangiates in comparison to the basal ferns [33], however, still maintained the IR structure. Thus, DR structure only occurred in the early diverging subg. *Rupestrae* (Fig. 4 a, b) and *S. tamariscina*. (Fig. 4 c).

**Fig. 4.**
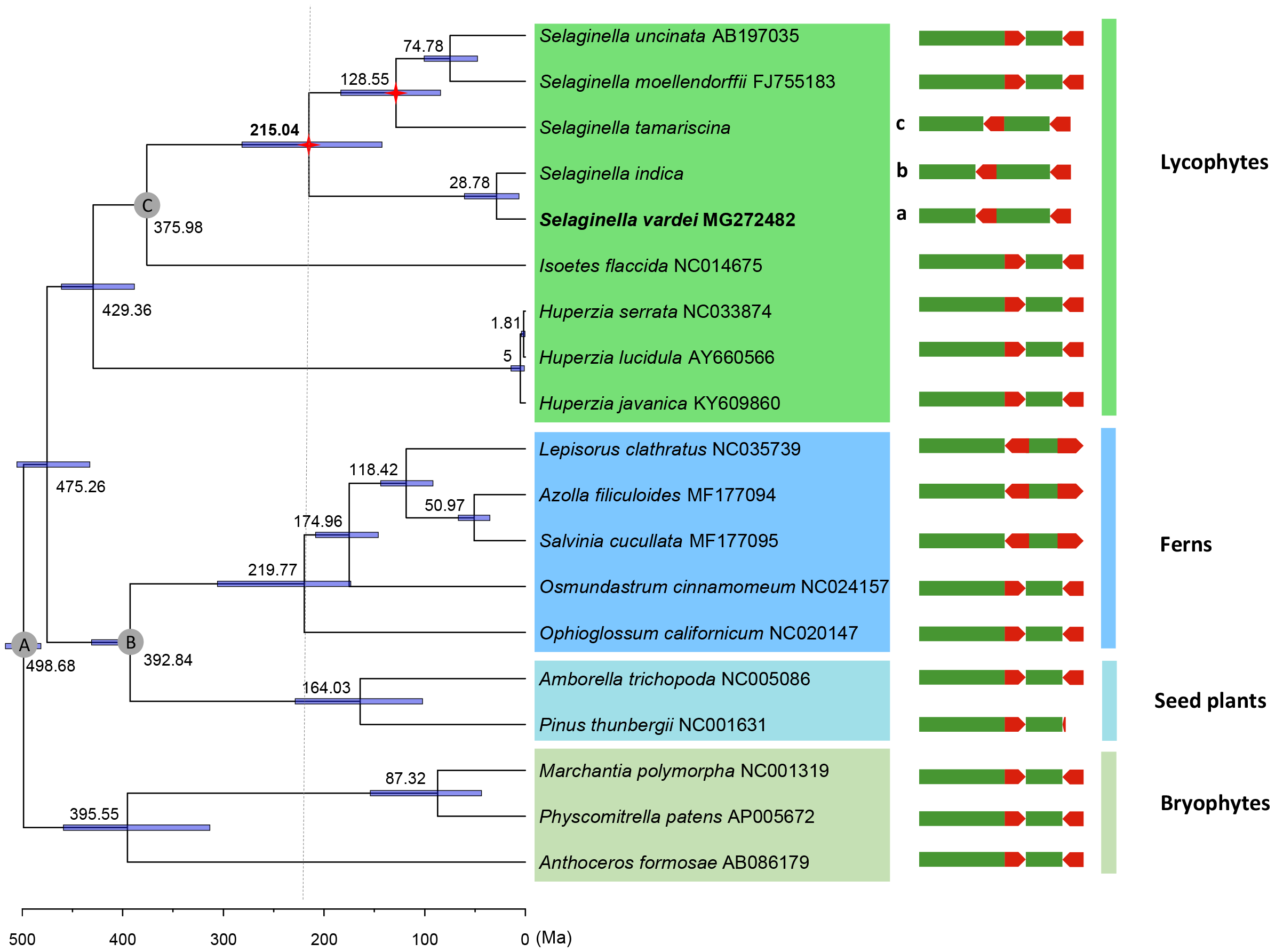
Phylogenetic reconstruction, time estimation of *S. vardei* and simplified structure of plastomes of each species. **Node A-C** represent the calibration nodes. Node A: a secondary calibration of the root age corresponded to the split of Bryophytes and vascular plants (470-583 Ma); node B: a fossil node separating ferns and seed plants (385-451 Ma); node C: a fossil node separating Selaginellaceae and its sister family Isoetaceae (372-392 Ma). **a, b, c** represents the plastomes with DR structure.

## Discussion

### The Characterization of the DR Structure in the Plastomes of Selaginellaceae

It is well known that plastome organization is highly conserved within land plants with the IR being the most conservative feature of land plant plastomes [3]. However, with the studies on plastomes going deeper, loss of one copy of the IR was uncovered to occur in several lineages of seed plants [6–8], and the DR structure has been reported in the recently published plastome of *S. tamariscina* with the mechanism of DR structure occurrence and maintenance left unknown [27]. Here we documented that the plastome of the *Selaginella* species *S. vardei* and *S. indica* possesses two copies of DR, rather than IR. The two copies of DR are identical and separate the single copy regions into almost equal size (Fig. 1). Furthermore, the DR structure of *S. vardei* was confirmed by PCR amplification in a different individual of *S. vardei* (collected from Tibet, China) and two other representative species (two individuals of *S. indica* from China and one individual of *S. dregei* from Kenya) (Table 1, Additional file 2, Fig. S2**a**), suggesting that the DR structure is a shared characteristic in subg. *Rupestrae. Selaginellaa* subg. *Rupestrae* containing *S. vardei* and *S. indica* is an early diverging clade, which is sister to *S. tamariscina* and the clade containing *S. moellendorffii* and *S. uncinata* (Fig. 4). Therefore, more representative samples of other subgenera are needed to elucidate whether plastomes with DR in subg. *Rupestrae* and *S. tamariscina* a synapomorphy or autopomorphy.

The DR structure, first reported in a red alga P. *purpurea* [26], originated independently in Selaginellaceae from red algae, since they are quite distantly related. Considering the ancestral plastome in lycophytes has the typical IR as shown in plastomes of the sister families, Lycopodiaceae [34–36] and Isoetaceae [37], we propose that the innovative DR structure of plastomes in subg. *Rupestrae* and *S. tamariscina* [27] originated within the family Selaginellaceae. Comparing with the plastome structure of *I. flaccida*, a species from its sister family Isoetaceae, we detect a notable inversion of a ca. 50-kb from *trn*N-*trn*F in *S. vardei*. The ca. 50-kb inversion starts from the LSC and terminates at the border of DRb/SSC, spanning one DR copy, thus, results in the orientation change from inverted to direct. According to our divergence time estimation, the DR structure of subg. *Rupestrae* occurred at late Triassic (ca. 215 Ma) with the 95% confidence interval as 142.5-281.6 Ma (Fig. 4). The previous studies inferred the divergence time of subg. *Rupestrae* around 200 Ma [32, 38], which lent support to our current results. The well-known Permian-Triassic (P-T) extinction event, occurred about 252 Ma ago [39] falls into the confidence interval of DR occurrence, therefore, the DR structure in Selaginellaceae possibly reflects a relic of the environmental upheaval during the extinction event related to increased anoxia, aridity, and UV-B radiation [39].

### Conformation of the Plastome with DR Structure of Selaginellaceae

Although mapped as circular molecules, which are in a quite low proportion, the plastomes of land plants displayed a great structural diversity with a mixture of monomers and head-tail concatemers of circular and linear molecules together with highly complex branched structures [40, 41]. The presence of two equimolar isomers (direction change of single copy regions), which was hypothesized to occur through flip-flop recombination of the two IR regions in a circular plastome [10, 42], has been recently proved to be the results of recombination-dependent replication (RDR) among linear plastome templates [43, 44]. However, the existence of direct repeats could promote multipartite chromosomal architecture as proposed in mitogenomes [45]. Repeats larger than 1 kb are typically at or close to the recombinational equilibrium as shown in the mitogenome of *Ginkgo biloba* L., the recombinant forms of two 5.3 and 4.1 kb repeats existing at roughly similar stoichiometries [46]. In plastome of *Monsonia* (Geraniaceae), the direct repeats (2-3 kb) were confirmed to produce alternative structures based on the RDR mechanism [44]. Thus, progresses in understanding the structure of both mitogenomes and plastomes suggest the existence of multipartite subgenomes for plastomes with DR in Selaginellaceae. The ca. 13 kb DR region in *S, vardei* could promote the generation of subgenomic chromosomes through recombination between two DR regions within one plastome or between different molecules following the RDR mechanism, with most or all of these molecules potentially occurring at similar stoichiometries. Two alternative read assemblies mapped at both ends of two copies of DR regions reflect the existence of subgenomes in plastome of *S. vardei* (Additional file 4, Fig. S4). Furthermore, we screened the PacBio reads of *S. tamariscina* plastome from Genbank and selected seven reads spanning the whole DR with both ends adjacent to genes from LSC, and two reads spanning DR with both ends adjacent to genes of SSC (Additional file 5, Fig. S5) confirming the recombination between DR regions and the existence of subgenomes in plastomes with DR structure. Either the master chromosome or subgenomic chromosomes could form head to tail multimers or branched complex based on the RDR mechanism (Additional file 4, Fig. S4**e**).

### The Co-occurrence of DR and Extremely Few SDR in the Plastome of Selaginellaceae

Remarkably, SDRs in plastomes with DR are extremely depauperate, as is shown in *S. indica* and *S. vardei* (Fig. 3e, f). There is no SDRs located at the ends of this ca. 50-kb inversion and the size of all the SDRs in *S. vardei* (six) and *S. indica* (five) is 16-25 bp, which only possibly invoke microhomology-mediated recombination. Repeats mediating homologous recombination in plastomes and mitogenomes appears to be limited to large repeats (>100-200 bp) [43]. Plastomes with DR could generate subgenomes at roughly similar stoichiometries by the recombination of DR regions. Therefore, the co-occurrence of DR and extremely few SDRs suggest that plastomes with DR are possible susceptive to SDRs scattered in single copy regions, which could generate secondary recombination within and among subgenomes [46, 47]. This secondary recombination caused by direct or inverted short repeats possibly results in plastome fragmentation or gene loss, which may influence the survival of plants. Therefore, we hypothesis that the co-occurrence of DR, together with virtually no other recombinational SDRs, maintains the integrity and stability of plastome, and survives long evolutionary history with over 400 million years.

## Conclusions

We documented the unconventional DR structure in plastome of *S. vardei* and two other representative species in subg. *Rupestrae sensu* Weststrand and Korall (2016). The DR structure occurred within the Selaginellaceae group and most likely arose at late Triassic (ca. 215 Ma). We propose that the ca. 50-kb inversion resulted in the DR structure, which could generate multipartite subgenomes induced by the recombination between DR regions. Subsequently, the co-occurrence of DR and few SDRs plays key role in maintaining the integrity and stability of the unconventional plastomes, and survives long evolutionary history with over 400 million years.

## Methods

### Taxon Sampling, DNA Extraction, Sequencing and Assembly

*Selaginella vardei* is a member of the monophyletic subgenus, subg. *Rupestrae sensu* Weststrand and Korall (2016), characterized by having monomorphic and helically arranged vegetative leaves and tetrastichous strobili. We collected a sample of *S. vardei (Zhang 6948*, PE) from the wile in Sichuan Province, China for this study and deposited a voucher of the collection in the Herbarium of Institute of Botany, CAS (PE). One closely related species, *S. indica*, was also collected from Yunnan Province and deposited in PE. **One of the authors, Prof. Xian-Chun Zhang, a proficient taxonomist on ferns and lycophytes, identified the samples. Both species are not endangered according the Convention on International Trade in Endangered Species of Wild Fauna and Flora (CITES)** (https://www.cites.org/).

Total genomic DNA was isolated from silica gel-dried materials with a modified cetyl-trimethylammonium bromide (CTAB) method [48]. Library construction was performed with the NEBNext DNA Library Prep Kit (New England Biolabs, Ipswich, Massachusetts, USA) and sequencing was performed on the Illumina HiSeq 2500 (Illumina, San Diego, California, USA). Illumina paired-end reads were mapped to *S. uncinata* (AB197035) [30] and *S. moellendorffii* (FJ755183) [31], with medium-low sensitivity in five to ten iterations in Geneious 9.1.4 (Biomatters, Inc., Auckland, New Zealand; https://www.geneious.com). The mapped reads were then assembled into contigs in Geneious. Additionally, the cleaned reads were assembled *de novo* with SPAdes v. 3.10.1 [49] using a range of kmer sizes from 21 to 99. Putative plastome contigs were identified using BLASTN 2.2.29 [50], with the previously published *S. uncinata* and *S. moellendorffii* plastomes as reference. We also used bandage v. 0.8.1 [29], a program for visualizing *de novo* assembly graphs, to help select plastome contigs and analyze *de novo* assembly results by importing the fastg file created by SPAdes. The contigs obtained from above ways were then combined and imported into Geneious to extend and assemble into the complete plastomes.

### Gene Annotation

Gene annotations were performed using local BLAST with default parameter settings [51]. Putative start and stop codons were defined based on similarity with known sequences. The tRNAs were further verified using tRNAscan-SE version 1.21 [52] and ARAGORN [53]. Circular and linear genome maps were drawn with OGDraw version 1.2 [54].

### PCR Confirmation of *S. vardei* Plastome

To confirm the accuracy of plastome assembly of *S. vardei*, we designed 12 pairs of primers (Additional file 7, Table S2) using Primer v 3.0 [55] based on the assembled sequence order of *S. vardei* to do long range PCR trying to amplify and sequence the whole plastome, and then, compare with the assembled one. Furthermore, we designed twelve pairs of primers (Additional file 9, Table S4) at the boundaries of the DR structure and the ca. 50-kb inversion in plastome of *S. vardei* (marked on Fig. 1) to confirm the accuracy of assembly. The PCR amplifications were performed in a total volume of 20 μL containing 4 μL of 5X PrimeSTAR GXL Buffer, 1.6 μL of dNTP Mixture (2.5 mM each), 1.2μL of each primer (5 mM), 0.4 μL of PrimeSTAR GXL DNA Polymerase and 20 ng of template DNA. Cycling conditions were 98 °C for 3 min, followed by 40 cycles of 98 °C for 10 s, 58 °C for 30 s and 72 °C for 5 min for long range PCR and 1.5 min for normal PCR, and a final extension of 72 °C for 10 min. The PCR products were verified by electrophoresis in 0.8% agarose gels stained with ethidium bromide. Then, we designed the internal primers (Additional file 8, Table S3) to get sequences of these PCR products using Sanger sequencing. The PCR products of the DR and inversion confirmation were also sequenced by the company Majorbio, Beijing, China.

### PCR Confirmation of DR Structure in Related Representatives of subg. *Rupestrae*

To further test whether the structure found in *S. vardei* existed in other species of subg. *Rupestrae*, PCR amplification using primers designed at the boundaries of the DR structure and the ca. 50-kb inversion in plastome of *S. vardei* (marked on Fig. 1, additional file 9, Table S4) were carried out with another individual of *S. vardei*, two individuals of *S. indica*, and one individual of *S. dregei* (Table 1). Only positive control was carried out in these species. The PCR procedure follows the conditions of normal PCR mentioned above. The PCR products were verified by electrophoresis in 0.8% agarose gels stained with ethidium bromide. The PCR products were sequenced by the company Majorbio, Beijing, China.

### Comparison of the Plastomes of *S. vardei* and Other Lycophytes

Dot plot analyses of the plastid genome of *S. vardei* and other lycophytes (*S. uncinata* AB197035, *S. moellendorffii* FJ755183, *I. flaccida* NC_014675 and *H. serrata* NC_033874) were performed using the Gepard Software [56] in order to identify the putative structural rearrangements in *S. vardei* plastome. The syntenic analyses of linear maps for plastomes of *S. vardei* and other lycophytes were then carried out based on the dot plot analyses. The first site of LSC at the border of LSC/IRa was considered as the starting point.

### Repeat Analyses

Short dispersed repeats (SDRs) were identified using RepeatsFinder [57] with default parameters. The circular layouts of SDRs in plastome were then visualized using the *circlize* package [58] in R. The accurate locations of some SDRs were marked in order to find some possible correlations with rearrangements. Six plastomes (*H. serrata* NC_033874, *I. flaccida* NC_014675, *S. uncinata* AB197035, *S. moellendorffii* FJ755183, *S. indica* and *S. vardei*) were included. One copy of the IR/DR was removed from all plastomes used.

### Phylogenetic Analyses and Divergence Time Estimation of DR Structure

Thirty-two conservative protein-coding genes of plastomes were performed to reconstruct the phylogenetic framework using 19 species from previously published plastomes of land plants (from moss to seed plants) (Additional file 11, Table S6). We downloaded the raw reads of *S. tamariscina* from Genbank (SRR6228814, SRR7135413), assembled plastid contigs and extracted the 32 gene sequences to perform the phylogenetic analysis since the complete plastome of it has not been released on the Genbank. A total of 19,248 bp sequences were aligned using MAFFT [59] under the automatic model selection option with some manual adjustments. Maximum-likelihood (ML) analysis was performed using RAxML v. 7.4.2 with 1000 bootstrap replicates and the GTR+G model [60] based on Akaike information criterion (AIC) in jModeltest 2.1.7 [61]. The simplified structure of the plastome of each species was then mapped on the phylogenetic tree showing the direction of rRNA encoding repeat.

Estimations of lineage divergence times were undertaken in BEAST version 1.8.2 [62] with three fossil calibration nodes were employed. A secondary calibration of the root age corresponded to the split of Bryophytes and vascular plants (Fig. 4, node A: [470‐583 Ma]) [63] with a selection of normal prior distributions. We employed two fossils for the second (separating Selaginellaceae and its sister family Isoetaceae (372-392 Ma) [32]) and third (separating ferns and seed plants (385-451 Ma) [63] nodes with a lognormal prior distribution. A relaxed clock with lognormal distribution of uncorrelated rate variation was specified. A birth-death speciation process with a random starting tree was adopted. The MCMC chain was run for 400,000,000 generations, sampling every 1000 generations. The effective sample size (ESS) was checked in Tracer v 1.5 [64]. The maximum clade credibility tree was generated using TreeAnnotator in BEAST and the tree was plotted using FigTree 1.4.3 [65].

## Abbreviations

CITES, Convention on the Trade in Endangered Species of Wild Fauna and Flora; CTAB, Cetyl-trimethylammonium bromide; DR, Direct repeat; IR, Inverted repeat; LSC, Large single copy; ML, Maximum likelihood; NGS, Next-generation sequencing; RDR, Recombination-dependent replication; rRNA, Ribosomal RNA; SDRs, Short dispersed repeats; SSC, Small single copy; tRNA, Transfer RNA

## Declarations

## Acknowledgments

We are grateful to Shi-Liang Zhou, Ran Wei, Shan-Shan Dong, Wen-Pan Dong and Jong-Soo Kang for helpful discussion. We thank Dr. Bing Liu for providing materials, Chang-Hao Li for help with analyses, and Li Wang, Zhi-Qiang Wu, AJ Harris for helpful revision and polishing of the whole manuscript. We also thank two anonymous reviewers for the insightful comments on the manuscript.

## Ethics approval and consent to participate

Not applicable.

## Consent for publication

Not applicable.

## Availability of data and materials

The newly sequenced plastome sequences have been deposited in Genbank under the accession number MG272482 and MK156801.

## Authors’ contributions

XCZ collected samples from the field, QPX and XCZ planned and designed the research, HRZ performed experiments and analyzed data, HRZ and QPX wrote the manuscript. All authors discussed the results and commented on the manuscript. All authors read and approved the final manuscript.

## Additional materials

**Additional file 1: Fig. S1. Confirmation of *S. vardei* plastome structure. a:** PCR amplification results of *S. vardei*. Primer pairs used are indicated at the top of each lane. Size markers are in bp. Gene names of these 11 products are as follows: 1: *rpo*B – *rps*2; 2: *rps*2 – *chl*B; 3: *chl*B – *ycf*2; 4: *ycf*2 – *psa*B; 5: *psa*B – *rrn*23; 6: *rrn*23-*rpl*2; 7: *rpl*2 – *pet*B; 8: *pet*B – *pet*E; 9: *pet*E – *atp*B; 10: *atp*B – *ycf*1; 11: *ycf*1-*rrn*23; 12: *rrn*23-*rpo*B. **b:** PCR confirmation results of DR structure and Inversion. Primer pairs used are indicated at the top of each lane. Size markers are in bp. Gene names of these 12 fragments are as follows: 1-2: *rps*4 – *rrn*5; 1-3: *rps*4 – *pet*N; 3-4: *pet*N – *rpl*2; 2-4: *rrn*5 – *rpl*2; 1-2: *rps*4 – *rrn*5; 1-5: *rps*4 – *atp*E; 5-6: *atp*E – *chl*L; 2-6: *rrn*5 – *chl*L; 7-8: *ccsA* – *rrn*5; 7-9: *ccsA* – *pet*N; 9-10: *pet*N – *rpo*B; 8-10: *rrn*5 – *rpo*B; **c:** Reads coverage of *S. vardei* plastomes; **d:** The assembled graph in bandage showing DR structure.

**Additional file 2: Fig. S2. Plastome structure confirmation and sequence alignments of related representatives. a:** PCR confirmation results of DR structure in another individual of *S. vardei* and two representative species within subg. *Rupestrae*. Primer pairs used are indicated at the top of each lane. Size markers are in bp. Gene names of these fragments are as follows: 1-2: *rps*4 – *rrn*5; 3-4: *pet*N – *rpl*2; 5-6: *atp*E – *chl*L; 7-8: *ccs*A – *rrn*5; 9-10: *pet*N – *rpo*B; **b:** Alignment results of sequences from each product in **a**, the light grey color represents regions with identical base pairs among individuals, whereas the dark color highlights regions with inconsistent base pairs.

**Additional file 3: Fig. S3. Dot-plot analyses of plastomes between *S. vardei* and other lycophyte species.**

**Additional file 4: Fig. S4. Reads coverages of putative subgenomes and alternative reads assemblies in *S. vardei*. a, b:** Two alternative reads assemblies at LSC/DRa and SSC/DRb boundaries: the unmatched reads in black box in **a** is consistent with assembled sequence in black box in **b**, the unmatched reads in red box in **b** is consistent with assembled sequence in red box in **a; c, d:** Two alternative reads assemblies at DRa/SSC and DRb/LSC boundaries: the unmatched reads in black box in **c** is consistent with assembled sequence in black box in **d**, the unmatched reads in red box in d is consistent with assembled sequence in red box in **c. e:** the simplified structure for master chromosomes and subgenomes based on a, b, c, d in this figure. We define the arrow end as B, and the other end as A. End B of either LSC (**a**) or SSC (**b**) can be assembled with end B of DR. End A of either LSC (**c**) or SSC (**d**) can be assembled with end A of DR.

**Additional file 5: Fig. S5. Screened PacBio reads of *S. tamariscina* showing evidence of the existence of subgenomes in plastomes with DR. a:** the simplified plastome structure of *S. tamariscina* based on Xu et al. (2018); **b, c:** the simplified subgenome structure of plastomes with DR, supported by the screened PacBio reads of *S. tamariscina* plastome as listed.

**Additional file 6:** Table S1. Genes present in the plastome of *S. vardei*.

**Additional file 7:** Table S2. Primers designed for long range PCR amplification of plastome of *S. vardei*.

**Additional file 8:** Table S3. Internal primers designed for Sanger sequencing of PCR products of *S. vardei*.

**Additional file 9:** Table S4. Primers designed for Sanger sequencing of PCR confirmation of *Selaginella* subg. *Rupestrae*.

**Additional file 10:** Table S5. Detailed dispersed short repeats in plastid genome of each species.

**Additional file 11:** Table S6. Species selected in the phylogenetic analyses.

