## Supplementary figures and images for "Direct Repeats Co-occur with Few Short Dispersed Repeats in Plastid Genome of A Spikemoss, *Selaginella vardei* (Selaginellaceae, Lycophyta)"

### Additional file 2, Fig. S1

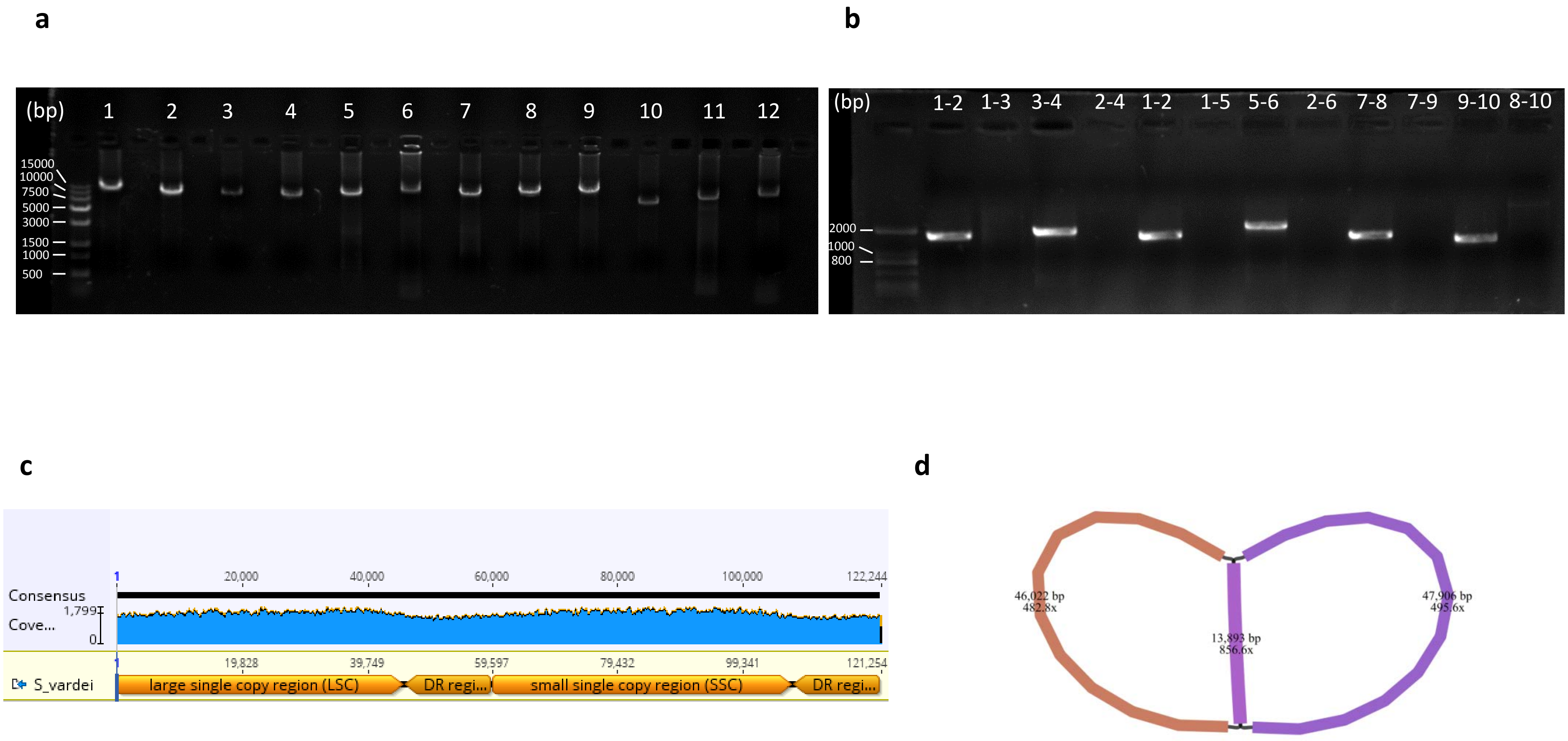

### Additional file 2, Fig. S2

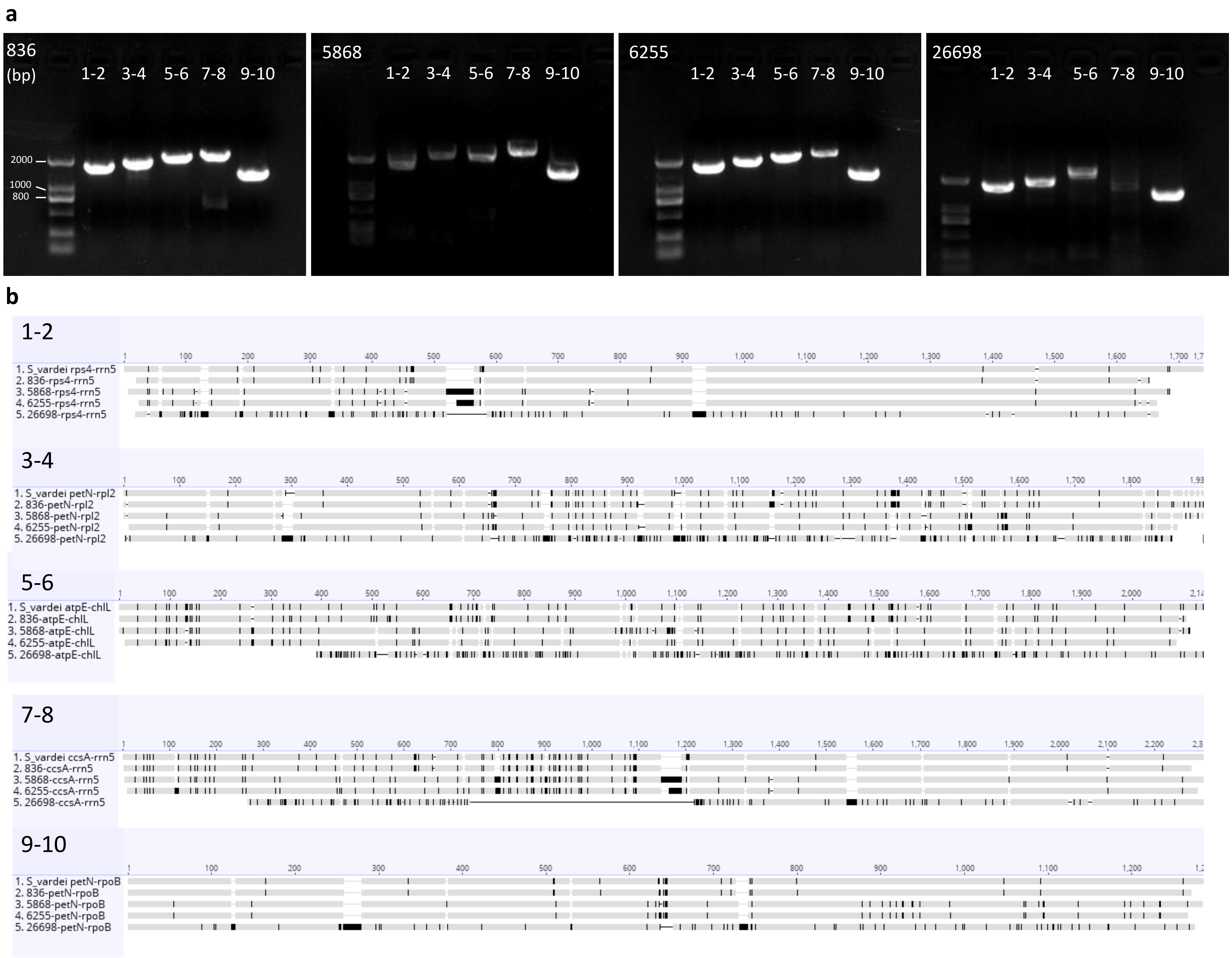

### Additional file 2, Fig. S3

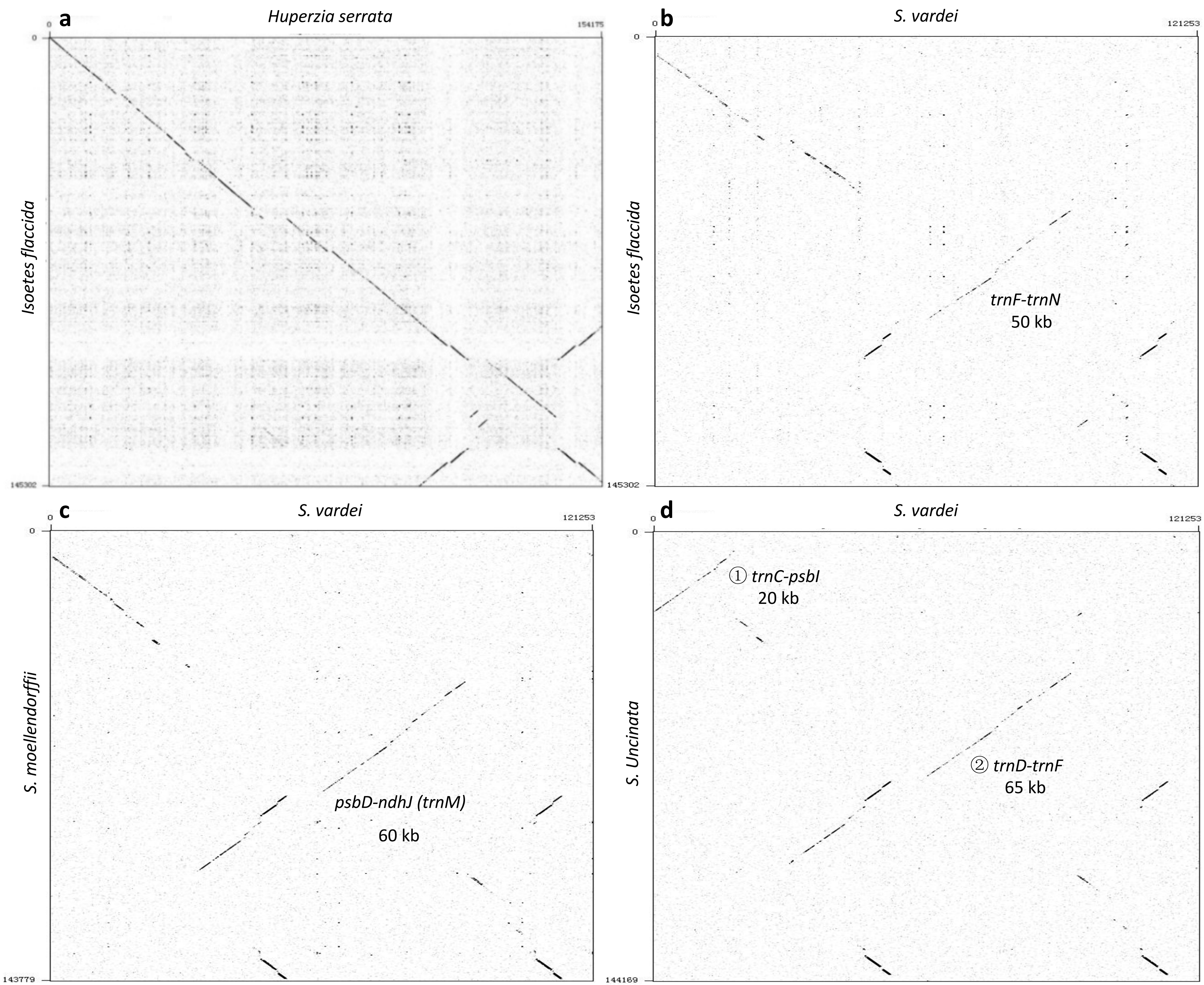

### Additional file 2, Fig. S4

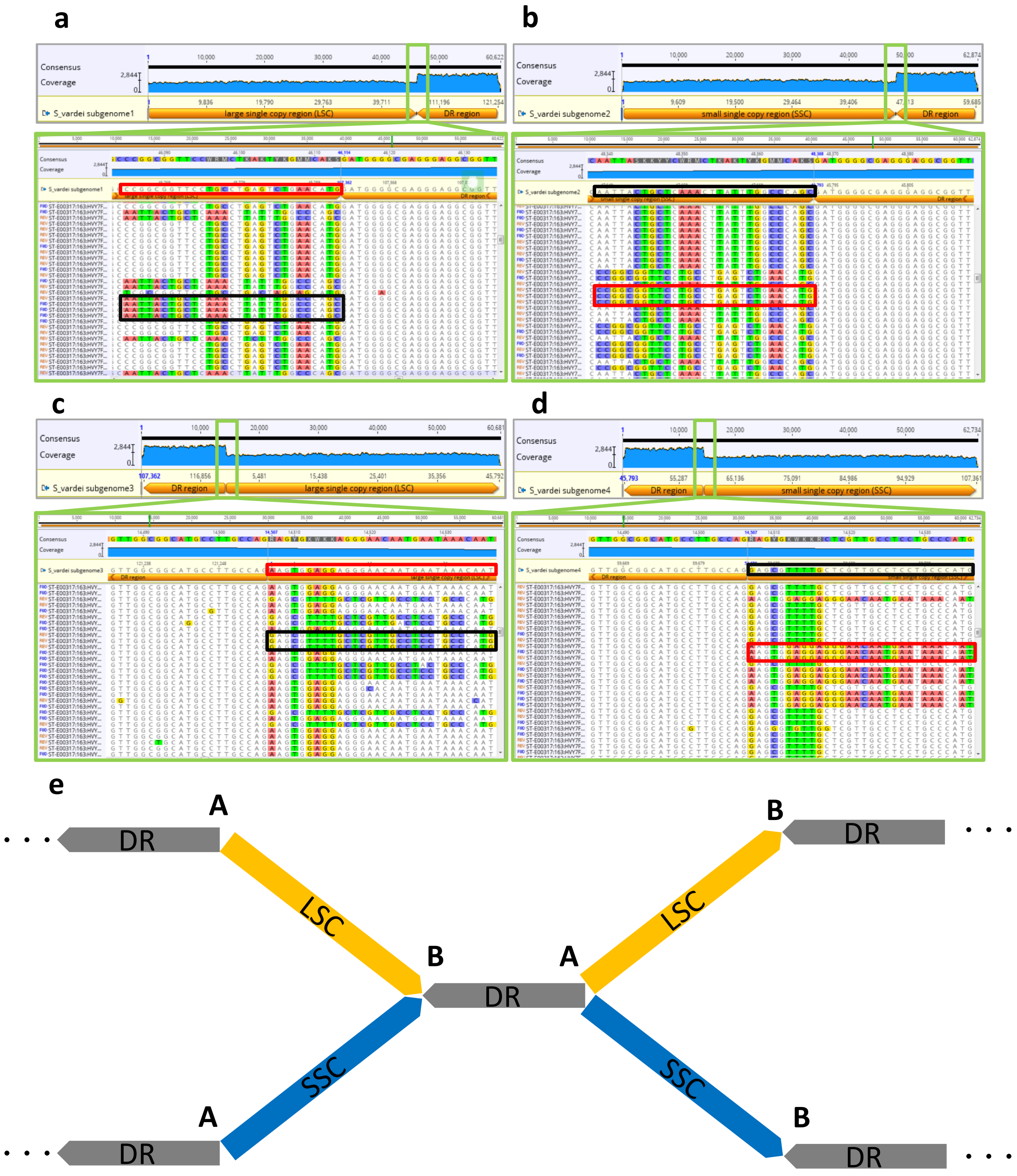

### Additional file 2, Fig. S5

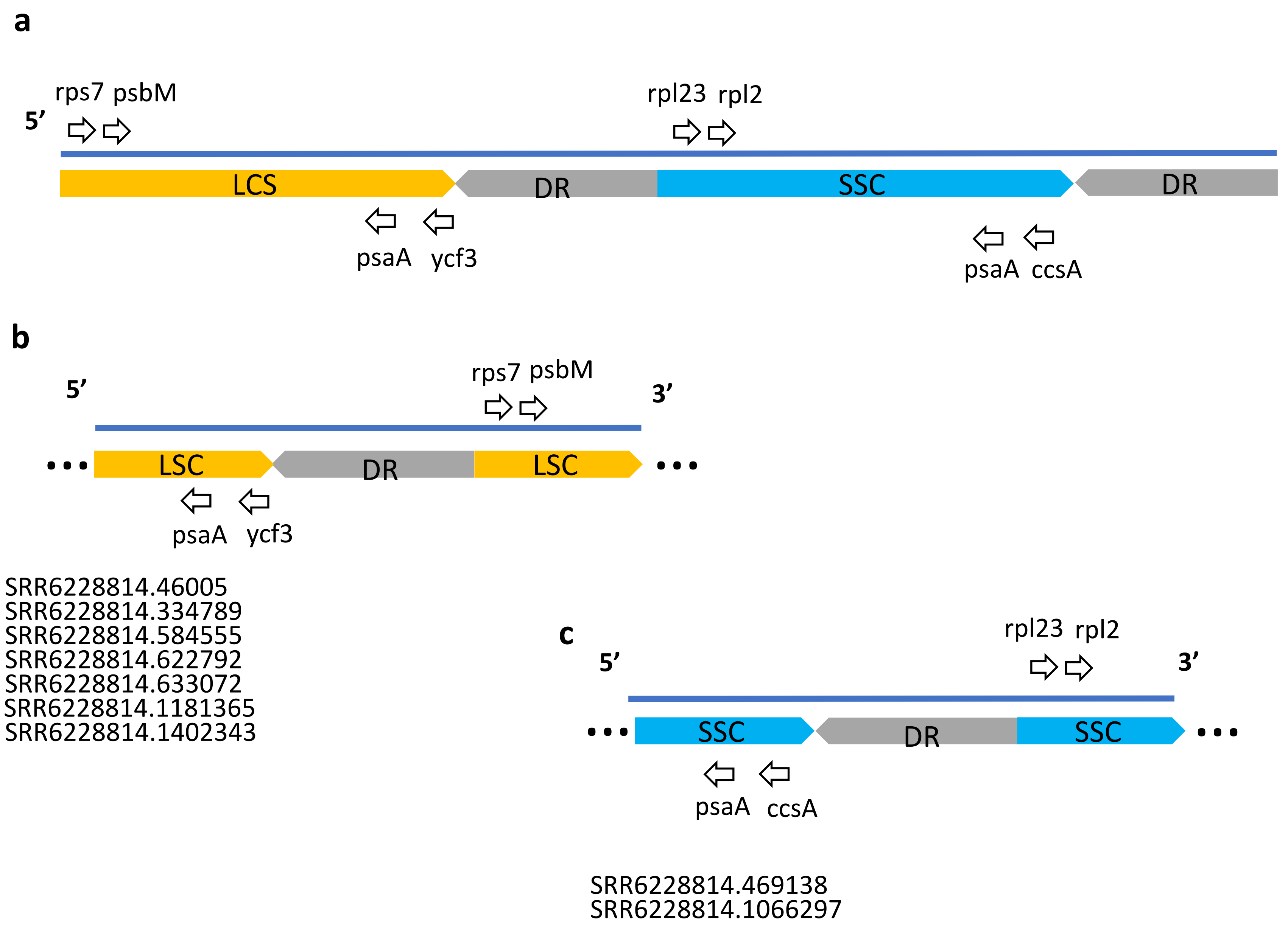
