## Supplemental Data 1 for "Direct Repeats Co-occur with Few Short Dispersed Repeats in Plastid Genome of A Spikemoss, *Selaginella vardei* (Selaginellaceae, Lycophyta)"

Table S1 Genes present in the plastome of *S. vardei*.

| **Chloroplast genome feature** | **Genes** |
| --- | --- |
| Photosystem I | *psaA, psaB, psaC, psaI, psaJ* |
| Photosystem II | *psbA, psbB, psbC, psbD, psbE, psbF, psbH, psbI, psbJ, psbK, psbL, psbM*^2^*, psbN, psbT, psbZ* |
| Cytochrome *b_6_ /f* | *petA, petB*^1^*, petD*^1^*, petG, petL, petN*^2^ |
| ATP synthase | *atpA, atpB, atpE, atpF, atpH, atpI* |
| RuBisCO | *rbcL* |
| Large subunit ribosomal proteins | *rpl2, rpl16, rpl20, rpl21, rpl22, rpl36* |
| Small subunit ribosomal proteins | *rps2, rps3, rps4, rps7*^2^*, rps8, rps11, rps14, rps18, rps19* |
| RNA polymerase | *rpoA, rpoB, rpoC1*^1^*, rpoC2* |
| Unknown function protein-coding gene | *ycf1, ycf2, ycf3*^1^*, ycf4, ycf12* |
| Other genes | *ccsA, chlB, chlL, chlN, clpP, infA* |
| Ribosomal RNAs | *rrn16*^2^*, rrn23*^2^*, rrn4.5*^2^*, rrn5*^2^ |
| Transfer RNAs | *trnC-GCA*^2^*, trnD-GUC, trnE-UUC, trnF-GAA, trnfM-CAU, trnH-GUG, trnM-CAU, trnN-GUU*^2^*, trnR-ACG, trnW-CCA, trnY-GUA* |

^1^ Genes containing a single intron.

^2^ Two gene copies in IRs.

Table S2. Primers designed for long range PCR amplification of plastome of *S. vardei*.

| **Primers** | **Sequences (5'to3')** |
| --- | --- |
| 1-*rpo*B-F | ATTTCCCAAGGATCGAGGATATGG |
| 1-*rps*2-R | CGAGCGGTTTGGGTTAGATCTATA |
| 2-*rps*2-F | CACTACGTGAATGAGAAATGGCTC |
| 2-*chl*B-R | CTGGAATGATTTGATTAACCCCGG |
| 3-*chl*B-F | CGAGGGTACTAATCAAATAGCGGA |
| 3-*ycf2*-R | TTAGCAATTGATTTATTGGGGGCC |
| 4-*ycf2*-F | ATGCCCCTACCTGCCATATTATTT |
| 4-*psa*B-R | CCGGTAGCTCTTTCCATAGTACAA |
| 5-*psa*B-F | ATTCAGCGTTCTTGAACCAGGATA |
| 5-*rrn*23-R | ACACAGGTGGGTAGGTAGAGAATA |
| 6-*rrn*23-F | ATCTCCGGATCCATGCTTATTTGT |
| 6-*rpl*2-R | TCGGGGTGCAAAATATAACCTTTG |
| 7-*rpl*2-F | CCTAATATCCCAAAACTGCCTTGC |
| 7-*pet*B-R | TTACCAGCAAATATGTCCCTCCTC |
| 8-*pet*B-F | GTGTTGACATGAGGAGGGACATAT |
| 8-*psbE*-R | TCATCGACTTGCTCCGACGAATTG |
| 9-*psbE*-F | ATCCATAGCATCACCATACCTTCC |
| 9-*atpB*-R | TTCATTTGACCATAAACCAAGGCC |
| 10-*atpB*-F | AGACCCCTTGGATTCAACTTCTAC |
| 10-*ycf*1-R | ACTAATAATGCCACTCAGGGGTTT |
| 11-*ycf*1-F | TTCGATTGTGGCTATACATAGGGG |
| 11-*rrn*23-R | CTAGGGATAACAGGCTGATCTTCC |
| 12-*rrn*23-F | ATCTCCGGATCCATGCTTATTTGT |
| 12-*rpoB*-R | CCATATCCTCGATCCTTGGGAAAT |

Table S3 Internal primers designed for Sanger sequencing of PCR products of *S. vardei*.

| Primers | Primer sequences |
| --- | --- |
| 1-1 F | ATACTATGCGCGTAGGAATGAACA |
| 1-2 F | AGGACTCCGTACTTATTAGCGAAC |
| 1-3 F | GAAATGCCCACTAGAAAATCCGAC |
| 1-4 F | AGATCATTCGTCCAGGGTTCATAG |
| 1-5 F | TAGGCGGGGGTAAAATGATTAGTT |
| 1-6 F | CCATGGATGATTTGCGGATAATGA |
| 1-7 F | GAATTGTGGATATTGCTACGCGAA |
| 1-8 F | GTCCCGACCCTTTATATGTTCGTT |
| 1-9 F | TCTCCCCTCGTACAATACTATCCA |
| 1-10 F | AGATACTATTTTCCAGGGCCTTCC |
| 2-1 F | GGGACCTAGCTAGAACTCAGATTG |
| 2-2 F | TATGAATGACTCTACCACTCGCAC |
| 2-3 F | TCTACGGATTAGTCGTAGCTTTGG |
| 2-4 F | GTTCAAGCGACGAATTACAATTGC |
| 2-5 F | TTACGACGATCTTACCAAACAAGC |
| 2-6 R | ATGAGTTCAGAAGTTATTGCGCAG |
| 2-7 R | CATACGGTGGTTATTTCCCTCGTT |
| 2-8 R | ATCACCCGAAGTTTATGCGATTTC |
| 2-9 F | TTGAGTATGCGGTATTCCCTAAGG |
| 3-1 F | TGGATGGTGGTTCTCAATTCACTT |
| 3-2 R | TGATGGCACCGGAGATAATATTGT |
| 3-3 R | AGTGTTTATCATGTGCCTCTGGTA |
| 3-4 F | ACAATATTATCTCCGGTGCCATCA |
| 3-5 F | CATCCGCCGGAAAGAGAGAG |
| 3-6 R | AGCTTGCCCCTATGGTTCAC |
| 3-7 F | TGCTCAATATCCGCCACGAG |
| 3-8 R | AATTGGCGAGTCCGGTCAAT |
| 3-9 R | GAGAAGGAGGAGGTGGAGGT |
| 3-10 R | CATATCCATGTGCGGTTCCATATG |
| 4-1 F | CGAAATTCAACCGGGTCCATTAAA |
| 4-2 F | TGAGCTTTTCCATCCACCGG |
| 4-3 F | CAAACCATGTCCTTGCCACG |
| 4-4 R | CCACAGTAGCGGCAGAGAAT |
| 4-5 F | GTTGCAACTTTCTCACCGCC |
| 4-6 R | AGGAAAGTGGAGGCTTGAGC |
| 4-7 F | AACACAGCTTATCCCAGTGAATCT |
| 4-8 R | GATACCCACCAATAAGACTAGGCC |
| 4-9 R | GAGATGGGAGATTAGCAGGGAATT |
| 4-10 R | CCGGTAGCTCTTTCCATAGTACAA |
| 5-1 F | TAGCCTTATCCGGTATCAAACGAG |
| 5-2 F | CTTAGCCATGCCTCGTAATTTGAG |
| 5-3 R | CGCGCTTAGAAACTGATCCTTATG |
| 5-4 R | TAAGACCTTTACAATCGTAGCGGA |
| 5-5 R | GGCGAGACAACTGGTTACTCATAA |
| 5-6 F | TTTCCTTGCCCTTCTTACGAGTAA |
| 5-7 R | GAACTCGGTGGTGAAACTCTACTG |
| 5-8 F | TTCCTCTACGACTTAGACACCAGA |
| 5-9 R | TGCAGCTGAGGCATCCTAAC |
| 6-1 F | CTATGGGGTATTAGCAGCCGTTTC |
| 6-2 F | ATTCGTTATCCATCCCACGTAGAG |
| 6-3 R | CCTGATAGGTCGATCCGCTCATAC |
| 6-4 R | ATGCGGGGATATTTACTTCTTCCG |
| 6-5 R | GAGTGTTCAAGCTCTGTCTGTAGT |
| 6-6 R | TAGTGACTGGATGGCGGATACATG |
| 6-7 R | TTCTCGTTCATCCCGGGCTAGATA |
| 6-8 R | GGATATAGTCAATACTGCTTGGGC |
| 6-9 F | GGAAATATAAATTGAGGCAGCCCA |
| 6-10 F | ATCCCACGCCTTACCACTTG |
| 7-1 R | GGATAAATCTTTGGTTGTGCGGAC |
| 7-2 R | GCTGCGCAAATAGGAATCTATCTG |
| 7-3 R | CTCTATTTGCCTAGACGTGATCCA |
| 7-4 F | ATACTCTATGAGATGGGCGGAGTA |
| 7-5 F | CCCTTGACAGTATGATAACCGCTA |
| 7-6 F | GATTTGTGAGAATTGCCGGCTAAT |
| 7-7 F | CAGGACATTGTATTGCCACCTTTC |
| 7-8 R | GTCCTAGATCCAGCTATGATTGGG |
| 7-9 F | AGGGTACTGTCAGTGGTCCC |
| 8-1 F | GTGTTGACATGAGGAGGGACATAT |
| 8-2 R | GAATAATCTTCTCCGCTACACCCT |
| 8-3 R | TATCTGAAGCTTGGTCCAGAATCC |
| 8-4 F | TGGACGGGTCATAAAGGGTATAAC |
| 8-5 F | CCTTAGTTTATTGGTAGCGGGGTA |
| 8-6 F | CAAGATTTATGCACAGAGAACGGG |
| 8-7 R | TGCCCAAACAAATGAATGGATTGA |
| 8-8 F | CCTCCGAAGCAGACTCATATTCTT |
| 8-9 R | GTCGTTACATGAGCCGACCTATAT |
| 8-10 F | CTGGAGTTGTCGGGGTCAAA |
| 9-1 F | GAAGGATTCCTCTGTGGCTGATAG |
| 9-2 R | AGGAGGAGCAACAATACAGTGTAC |
| 9-3 F | TACACTGTATTGTTGCTCCTCCTT |
| 9-4 F | AATTGGGGGAGGTATACGATTCAG |
| 9-5 F | TAGTTGTCCCGATCCGAGTTAATC |
| 9-6 F | CCATAAGTAAGAGCCGGAAAACAC |
| 9-7 F | GGATAATATTTTGCCGAACCTCCC |
| 9-8 R | TCACTTTAGGTTTCGTGGATTTGC |
| 9-9 R | CTCCCGACTACGTGACCAAG |
| 9-10 F | AGTCATTCGGAATGCTGCCA |
| 10-1 F | GCGTGATGACCCCTAACCAA |
| 10-2 F | TCCGTCGGCCCATTAATGAG |
| 10-3 F | TTACGAGTCAAGGCCTGTGC |
| 10-4 R | TCCGCGGAAATCATGACTCC |
| 10-5 F | GTCATTCCCCACAAAACGCC |
| 10-6 F | ATTGTTCATGAGGTTGGTATCCCA |
| 11-1 F | TGGGGATACTGGAGTCCCTG |
| 11-2 F | GAGGGGCCACTCACTGATAC |
| 11-3 R | AACCCTGGCGAACTGATTCC |
| 11-4 F | TGTCTACCGGTTTCACCACG |
| 11-5 F | CGGACGGCTGGTAATCATGA |
| 11-6 F | GAAAACTGCAGTTACCCCGC |
| 11-7 F | CCCAGCCCGTCCATTTGTAT |
| 11-8 R | CTAGGGATAACAGGCTGATCTTCC |
| 11-9 F | TGTGCAAAGGGATAGGGATGTTAA |
| 11-10 R | TTAACATCCCTATCCCTTTGCACA |
| 11-11F | ATGTAAAAGGTGTGAATCCGCTTG |
| 12-1 R | TACCCCAGTAGTCCTAGCCG |
| 12-2 F | ATTAGCAGCCGTTTCCAGCT |
| 12-3 R | AACACATGCAAGTCGTACGG |
| 12-4 R | ATGGCGCTACAGGGAATTCC |
| 12-5 F | TGGAATCCGCACGAGGAAAA |
| 12-6 F | ACGGCATCTCTCATCGTTCC |
| 12-7 R | TCGTTCCTTTGCGATTTGGC |
| 12-8 F | GGGTATTATGGGCGATCGCA |
| 12-9 R | TTGTGTTCACAAGCTAGCGA |
| 12-10 R | TCAATACTGCTTGGGCTGCC |
| 12-11 R | TACCACCCTCTGGCAATGTG |

Table S4. Primers designed for PCR confirmation of *Selaginella* subg. *Rupestrae*.

| **Primers** | **Sequences (5'to3')** |
| --- | --- |
| 1-*rps4*-F | GGGCATATAACCAGGCTTT |
| 2-*rrn*5-R | GTAGAGGAACCACACCAATC |
| 3-*pet*N-F | GCAGCCCAAGCAGTATTGAC |
| 4-*rpl*2-R | TTGTCACCGTCTTCATAGT |
| 5-*atp*E-F | AACAAAGCTGCGGATCTCGA |
| 6-*chl*L-R | TTGATTTACCTGTGCCGCCT |
| 7-*ccsA*-6F | GAAAACTGCAGTTACCCCGC |
| 8-*rrn*5-R | GTAGAGGAACCACACCAATC |
| 9-*pet*N-F | GCAGCCCAAGCAGTATTGAC |
| 10-*rpoB*-R | GTTTCTCCCATCGGTTTCCAATTT |

Table S5-1 Detailed dispersed short repeats in cp genome of *Huperzia serrata.*

| q_start | q_end | length | s_start | s_end | length | mismatch | q_location | s_location |
| --- | --- | --- | --- | --- | --- | --- | --- | --- |
| 2768 | 2784 | 17 | 7406 | 7422 | 17 | 0 | *rpl22* | *rps11* |
| 7451 | 7468 | 18 | 79118 | 79101 | -18 | 0 | *rps11* | *atpA* |
| 10278 | 10294 | 17 | 40757 | 40773 | 17 | 0 | *petD-petB* | *trnF-GAA-trnL-UAA* |
| 11926 | 11947 | 22 | 89620 | 89597 | -24 | 2 | *petB-psbH* | *rpoC2* |
| 22289 | 22308 | 20 | 116350 | 116331 | -20 | 0 | *trnW-CCA* | *trnN-GUU* |
| 23710 | 23726 | 17 | 78466 | 78450 | -17 | 0 | *petL-psbE* | *ycf12-trnR-UCU* |
| 30154 | 30172 | 19 | 30219 | 30201 | -19 | 0 | *ycf4-psaI* | *ycf4-psaI* |
| 30642 | 30660 | 19 | 105387 | 105405 | 19 | 0 | *psaI-accD* | *rps7-trnV-GAC* |
| 37024 | 37040 | 17 | 89012 | 88996 | -17 | 0 | *trnV-UAC* | *rpoC2* |
| 39699 | 39719 | 21 | 85693 | 85713 | 21 | 0 | *ndhJ-trnF-GAA* | *rpoC2* |
| 39997 | 40015 | 19 | 109837 | 109819 | -19 | 0 | *trnF-GAA* | *trnA-UGC* |
| 44574 | 44591 | 18 | 1329 | 1312 | -18 | 0 | *ycf3* | *rpl2* |
| 51797 | 51814 | 18 | 118683 | 118700 | 18 | 0 | *rps14* | *trnN-GUU-chlL* |
| 52430 | 52456 | 27 | 77336 | 77310 | -27 | 1 | *trnG-UCC* | *ycf12-trnR-UCU* |
| 74900 | 74917 | 18 | 5211 | 5194 | -18 | 0 | *psbK-psbI* | *rpl16-rpl14* |
| 74926 | 74961 | 36 | 74961 | 74926 | -36 | 0 | *psbK-psbI* | *psbK-psbI* |
| 77726 | 77746 | 21 | 40347 | 40366 | 20 | 1 | *ycf12-trnR-UCU* | *trnF-GAA-trnL-UAA* |
| 77773 | 77789 | 17 | 138567 | 138551 | -17 | 0 | *ycf12-trnR-UCU* | *rpl21-ndhF* |
| 84298 | 84321 | 24 | 84321 | 84298 | -24 | 0 | *atpI-rps2* | *atpI-rps2* |
| 85292 | 85315 | 24 | 85577 | 85600 | 24 | 1 | *rpoC2* | *rpoC2* |
| 85693 | 85713 | 21 | 39699 | 39719 | 21 | 0 | *rpoC2* | *ndhJ-trnF-GAA* |
| 87843 | 87878 | 36 | 87771 | 87806 | 36 | 2 | *rpoC2* | *rpoC2* |
| 97613 | 97655 | 43 | 97655 | 97613 | -43 | 1 | *petN-ycf66* | *petN-ycf66* |
| 103608 | 103640 | 33 | 103640 | 103608 | -33 | 4 | *rps7-trnV-GAC* | *rps7-trnV-GAC* |
| 103621 | 103647 | 27 | 127947 | 127972 | 26 | 3 | *rps7-trnV-GAC* | *rps15-ndhH* |
| 103987 | 104080 | 94 | 94 | 1 | -94 | 0 | *rps7-trnV-GAC* | *rps12-trnI* |
| 108722 | 108739 | 18 | 116387 | 116370 | -18 | 0 | *trnI-GAU* | *trnN-GUU* |
| 111360 | 111385 | 26 | 111479 | 111504 | 26 | 1 | *trnA-UGC-rrn23* | *trnA-UGC-rrn23* |
| 111422 | 111459 | 38 | 111607 | 111644 | 38 | 3 | *trnA-UGC-rrn23* | *trnA-UGC-rrn23* |
| 116549 | 116578 | 30 | 116595 | 116624 | 30 | 0 | *trnN-GUU-chlL* | *trnN-GUU-chlL* |
| 132665 | 132682 | 18 | 132682 | 132665 | -18 | 0 | *ndhG-ndhE* | *ndhG-ndhE* |

Table S5-2 Detailed dispersed short repeats in cp genome of *Isoetes flaccida*.

| q_start | q_end | length | s_start | s_end | length | mismatch | q_location | s_location |
| --- | --- | --- | --- | --- | --- | --- | --- | --- |
| 6875 | 6894 | 20 | 109935 | 109955 | 21 | 1 | *rpl36-rps11* | *ycf1-rps15* |
| 12322 | 12365 | 44 | 12365 | 12322 | -44 | 0 | *psbT-psbB* | *psbT-psbB* |
| 13872 | 13891 | 20 | 67133 | 67114 | -20 | 1 | *psbB* | *psbI* |
| 29383 | 29422 | 40 | 29422 | 29383 | -40 | 0 | *psaI-accD* | *psaI-accD* |
| 30087 | 30115 | 29 | 30115 | 30087 | -29 | 4 | *accD* | *accD* |
| 33215 | 33236 | 22 | 73587 | 73565 | -23 | 1 | *rbcL-atpB* | *atpH-atpI* |
| 33275 | 33291 | 17 | 108651 | 108635 | -17 | 0 | *rbcL-atpB* | *ycf1* |
| 35711 | 35728 | 18 | 104015 | 103998 | -18 | 0 | *trnV-UAC* | *trnN-GUU* |
| 36429 | 36446 | 18 | 36446 | 36429 | -18 | 0 | *trnV-UAC-ndhC* | *trnV-UAC-ndhC* |
| 45433 | 45456 | 24 | 45456 | 45433 | -24 | 2 | *ycf3-psaA* | *ycf3-psaA* |
| 57698 | 57714 | 17 | 128977 | 128961 | -17 | 0 | *trnT-GGU-trnE-UUC* | *ycf2* |
| 62364 | 62383 | 20 | 123567 | 123548 | -20 | 1 | *matK-rps16* | *ndhF-chlN* |
| 67205 | 67226 | 22 | 52014 | 51993 | -22 | 0 | *trnS-GCU* | *trnS-UGA* |
| 67206 | 67229 | 24 | 42352 | 42375 | 24 | 0 | *trnS-GCU* | *trnS-GGA* |
| 68975 | 69004 | 30 | 51204 | 51176 | -29 | 1 | *trnG-UCC* | *trnG-GCC* |
| 77326 | 77342 | 17 | 108206 | 108222 | 17 | 0 | *rpoC2* | *ycf1* |
| 81780 | 81796 | 17 | 62984 | 63000 | 17 | 0 | *rpoC1* | *matK-rps16* |
| 86505 | 86522 | 18 | 66093 | 66076 | -18 | 0 | *trnC-GCA* | *trnQ-UUG* |
| 89618 | 89726 | 109 | 75425 | 75533 | 109 | 2 | *ndhB* | *rps2-rpoC2* |
| 93414 | 93451 | 38 | 43907 | 43870 | -38 | 2 | *rps7-trnV-GAC* | *ycf3* |
| 97714 | 97739 | 26 | 35705 | 35729 | 25 | 1 | *trnI-GAU* | *trnV-UAC* |
| 98733 | 98751 | 19 | 39447 | 39429 | -19 | 0 | *trnA-UGC* | *trnF-GAA* |
| 103959 | 103978 | 20 | 20717 | 20698 | -20 | 0 | *trnN-GUU* | *trnW-CCA* |
| 103998 | 104015 | 18 | 97738 | 97721 | -18 | 0 | *trnN-GUU* | *trnI-GAU* |
| 123280 | 123297 | 18 | 74455 | 74472 | 18 | 0 | *ndhF-chlN* | *atpI-rps2* |
| 129285 | 129326 | 42 | 129326 | 129285 | -42 | 2 | *ycf2* | *ycf2* |
| 129626 | 129643 | 18 | 1515 | 1498 | -18 | 0 | *ycf2* | *rpl2* |
| 131308 | 131329 | 22 | 50943 | 50964 | 22 | 1 | *ycf2* | *rps14-trnfM-CAU* |

Table S5-3 Detailed dispersed short repeats in cp genome of *Selaginella uncinata*.

| q_start | q_end | length | s_start | s_end | length | mismatch | q_location | s_location |
| --- | --- | --- | --- | --- | --- | --- | --- | --- |
| 6158 | 6203 | 46 | 26015 | 25970 | -46 | 3 | *psbI* | *trnC-psbK* |
| 8406 | 8423 | 18 | 8423 | 8406 | -18 | 0 | *ycf12-atpA* | *ycf12-atpA* |
| 11484 | 11500 | 17 | 41714 | 41730 | 17 | 0 | *atpA-atpH* | *ycf2-trnF* |
| 11485 | 11501 | 17 | 41711 | 41727 | 17 | 0 | *atpA-atpH* | *ycf2-trnF* |
| 14941 | 14958 | 18 | 14958 | 14941 | -18 | 0 | *rpoC2* | *rpoC2* |
| 28374 | 28394 | 21 | 25744 | 25764 | 21 | 1 | *trnC-chlB* | *trnC* |
| 30693 | 30712 | 20 | 30712 | 30693 | -20 | 0 | *chlB-matK* | *chlB-matK* |
| 35005 | 35090 | 86 | 88502 | 88586 | 85 | 2 | *trnH* | *trnN-rps4* |
| 54612 | 54709 | 98 | 63898 | 63801 | -98 | 2 | *petA-psbJ* | *rpl20-psbB* |
| 54695 | 54711 | 17 | 4530 | 4514 | -17 | 0 | *petA-psbJ* | *rps7-psbM* |
| 58325 | 58342 | 18 | 88188 | 88205 | 18 | 0 | *trnW* | *trnN* |
| 65309 | 65329 | 21 | 72919 | 72898 | -22 | 1 | *psbB* | *rps8-rpl14* |
| 81912 | 81929 | 18 | 88244 | 88227 | -18 | 0 | *rrn16-rrn23* | *trnN* |
| 91822 | 91838 | 17 | 92191 | 92207 | 17 | 0 | *ndhF* | *ndhF* |
| 99531 | 99547 | 17 | 48270 | 48254 | -17 | 0 | *ndhI* | *atpB-rbcL* |
| 113402 | 113422 | 21 | 25744 | 25764 | 21 | 1 | *trnQ* | *trnC* |
| 116057 | 116075 | 19 | 82196 | 82178 | -19 | 0 | *trnD-trnY* | *rrn16-rrn23* |
| 120463 | 120486 | 24 | 94431 | 94408 | -24 | 1 | *psbC* | *rpl21-ccsA* |
| 126222 | 126239 | 18 | 126239 | 126222 | -18 | 0 | *psaA* | *psaA* |

Table S5-4 Detailed dispersed short repeats in cp genome of *Selaginella moellendorffii*.

| q_start | q_end | length | s_start | s_end | length | mismatch | q_location | s_location |
| --- | --- | --- | --- | --- | --- | --- | --- | --- |
| 7664 | 7685 | 22 | 28762 | 28741 | -22 | 1 | *trnC* | *trnQ* |
| 17320 | 17345 | 26 | 90940 | 90916 | -25 | 2 | *rpoC2* | *rrn5-trnR* |
| 19177 | 19196 | 20 | 4972 | 4953 | -20 | 0 | *rpoC2-atpI* | *ndhB-psbM* |
| 30714 | 30735 | 22 | 4762 | 4784 | 23 | 1 | *chlB-matK* | *ndhB-psbM* |
| 30715 | 30739 | 25 | 4759 | 4785 | 27 | 2 | *chlB-matK* | *ndhB-psbM* |
| 42960 | 43003 | 44 | 43003 | 42960 | -44 | 2 | *trnD-trnY* | *trnD-trnY* |
| 54375 | 54424 | 50 | 54424 | 54375 | -50 | 0 | *accD* | *accD* |
| 59643 | 59659 | 17 | 93831 | 93815 | -17 | 0 | *petA-psbJ* | *trnN-rps4* |
| 63725 | 63750 | 26 | 6245 | 6270 | 26 | 2 | *trnW-psaJ* | *psbM-petN* |
| 65911 | 65938 | 28 | 65880 | 65907 | 28 | 1 | *rpl20-clpP* | *rpl20-clpP* |
| 90669 | 90688 | 20 | 90640 | 90621 | -20 | 0 | *rrn4.5-rrn5* | *rrn4.5-rrn5* |
| 91729 | 91747 | 19 | 86464 | 86482 | 19 | 0 | *trnR-trnN* | *rrn16-rrn23* |
| 93447 | 93493 | 47 | 36147 | 36193 | 47 | 1 | *trnN-rps4* | *trnH* |
| 95235 | 95253 | 19 | 95180 | 95162 | -19 | 0 | *rps4-ndhF* | *rps4-ndhF* |
| 99187 | 99213 | 27 | 99160 | 99186 | 27 | 1 | *trnL-ccsA* | *trnL-ccsA* |
| 117634 | 117657 | 24 | 77681 | 77704 | 24 | 0 | *chlL-trnF* | *rps8-rpl14* |
| 129587 | 129603 | 17 | 130660 | 130676 | 17 | 0 | *ycf3* | *ycf3* |
| 129974 | 130015 | 42 | 130015 | 129974 | -42 | 0 | *ycf3* | *ycf3* |

Table S5-5 Detailed dispersed short repeats in cp genome of *Selaginella vardei*.

| q_start | q_end | length | s_start | s_end | length | mismatch | q_location | s_location |
| --- | --- | --- | --- | --- | --- | --- | --- | --- |
| 17397 | 17412 | 16 | 42152 | 42137 | -16 | 0 | *ycf12-psbI* | *psaA* |
| 25341 | 25356 | 16 | 58272 | 58257 | -16 | 0 | *ycf2* | *psbM-petN* |
| 25733 | 25748 | 16 | 33858 | 33843 | -16 | 0 | *ycf2* | *psbD* |
| 57908 | 57925 | 18 | 57974 | 57991 | 18 | 0 | *psbM-petN* | *psbM-petN* |
| 73582 | 73597 | 16 | 75269 | 75284 | 16 | 0 | *psbB* | *psbB-clpP* |
| 88082 | 88098 | 17 | 102166 | 102182 | 17 | 0 | *rbcL* | *ycf1-psaC* |

Table S5-6 Detailed dispersed short repeats in cp genome of *Selaginella indica*.

| q_start | q_end | length | s_start | s_end | length | mismatch | q_location | s_location |
| --- | --- | --- | --- | --- | --- | --- | --- | --- |
| 16906 | 16929 | 24 | 64579 | 64556 | -24 | 1 | *ycf12*-*psbI* | *rpl16* |
| 17404 | 17419 | 16 | 42074 | 42059 | -16 | 0 | *ycf12*-*psbI* | *psaA* |
| 17492 | 17508 | 17 | 60223 | 60207 | -17 | 0 | *ycf12*-*psbI* | *trnC*-*rpl2* |
| 46525 | 46544 | 20 | 79761 | 79780 | 20 | 0 | *trnN* | *trnW* |
| 88962 | 88978 | 17 | 103023 | 103039 | 17 | 0 | *rbcL* | *ycf1*-*psaC* |

Table S6 Species selected in the phylogenetic analyses

| **Species** | **Genbank No.** |
| --- | --- |
| *Amborella trichopoda* | NC 005086 |
| *Anthoceros formosae* | AB086179 |
| *Azolla filiculoides* | MF177094 |
| *Isoetes flaccida* | NC 014675 |
| *Huperzia javanica* | KY609860 |
| *Huperzia lucidula* | AY660566 |
| *Huperzia serrata* | NC 033874 |
| *Lepisorus clathratus* | NC035739 |
| *Marchantia polymorpha* | NC 001319 |
| *Osmundastrum cinnamomeum* | KF225592 |
| *Ophioglossum californicum* | NC 020147 |
| *Physcomitrella patens* | AP005672 |
| *Pinus thunbergii* | NC 001631 |
| *Physcomitrella patens* | AP005672 |
| *Salvinia cucullata* | MF177095 |
| ***Selaginella indica*** | ***MK156801** |
| *Selaginella moellendorffii* | FJ755183 |
| *Selaginella uncinata* | AB197035 |
| *Selaginella tamariscina* | - |
| ***Selaginella vardei*** | ***MG272482** |

- Genbank number unavailable

* newly sequenced plastome
